# Histone deacetylase 3 regulates microglial function through histone deacetylation

**DOI:** 10.1101/2022.09.08.507183

**Authors:** Laura Meleady, Morgan Towriss, Jennifer Kim, Vince Bacarac, Megan Rowland, Annie Vogel Ciernia

**Author notes:** **Correspondence:** Annie Vogel Ciernia.

## Abstract

**Background:** As the primary innate immune cells of the brain microglia respond to damage and disease through pro-inflammatory release of cytokines and neuroinflammatory molecules. Histone acetylation is an activating transcriptional mark that regulates gene expression, which is altered in states of disease. Inhibition of histone deacetylase 3 (Hdac3) has been utilized in pre-clinical models of disease to dampen inflammation, but the molecular mechanisms underlying Hdac3’s regulation of inflammatory gene expression in microglia is not well understood.

**Methods:** Functional changes in immortalized microglia were characterized using a Hdac3 specific inhibitor RGFP966 in response to an immune challenge lipopolysaccharide (LPS). Flow cytometry and cleavage under tags & release using nucleases (CUT & RUN) were used to investigate global and promoter-specific histone acetylation changes, resulting in altered gene expression.

**Results:** Hdac3 inhibition enhanced neuroprotective functions of microglia in response to LPS through reduced nitric oxide release and increased baseline phagocytosis. Inhibition of Hdac3 enhanced histone acetylation globally and at specific gene loci, resulting in the release of gene repression at baseline and enhanced responses to LPS.

**Conclusion:** The findings suggest Hdac3 serves as a negative regulator of microglial gene expression, and that inhibition of Hdac3 facilitates the microglial response to inflammation and its subsequent resolution. Together, this work provides new mechanistic insights into therapeutic applications of Hdac3 inhibition which mediate reduced neuroinflammatory insults through microglial response.

## Background

As the resident immune cells of the brain, microglia are acutely sensitive and respond rapidly to changes in the local brain environment^1–3^. Due to their ability to respond to a diverse number of stimuli, microglia are involved in virtually all CNS disorders, ranging from degenerative and neurodevelopmental diseases to autoimmune neuroinflammatory conditions ^4^. Dozens of genetic loci affecting microglial phagocytosis, activation, or immunoregulation have been linked to Parkinson’s disease (e.g., TREM2)^5^, Alzheimer disease (e.g., ABCA7, EPHA1, MS4A6A, CD2AP, CD33)^6^, frontotemporal dementia (e.g., GRN)^7^, schizophrenia (e.g., C4)^8^, and multiple sclerosis (MS) (e.g., TNFRSF1A, IRF8, CD6)^9–11^. Both over and under active microglial phenotypes have been linked to disease pathogenesis across different brain disorders ^4^. However, it is often unclear if altered microglial activity is helpful or harmful ^4^, necessitating a deeper understanding of microglial regulation and functional impacts on the brain in both health and disease.

In reaction to an immune insult, microglia rapidly increase expression of inflammatory cytokines, allowing for increased phagocytosis of infectious agents^12^. This requires induction of gene expression that is controlled through modifications of chromatin structure via epigenetic mechanisms^13^. Histone acetylation is an epigenetic regulator of active transcription, both at promoters (H3K9ac) and enhancers (H3K27ac). These acetylation marks promote more open chromatin by loosening the interactions between the DNA and histones and are recognized by transcriptional activators^14^. Histone acetylation is added by histone acetyltransferases (Hats) and removed by Histone deacetylases (Hdacs). Removal of histone acetylation alters DNA-chromatin electrostatic contacts resulting in compaction of chromatin structure, decreased accessibility for transcription factor binding and inhibition of transcription^14^.

Previous work has explored Hdac3 as a key negative regulator of gene expression in the brain^15^. Hdac3 is the only Hdac found in the N-CoR/SMRT complex^16^ and serves as the catalytic component of the complex, leading to histone deacetylation and transcriptional repression. In response to brain damage, translation of Hdac3 is upregulated in both cortical^23^ and spinal cord microglia^24^. The increase in Hdac3 protein levels appears to confer pro-inflammatory functions^23,24^ as conditional deletion of Hdac3 in microglia improves outcomes following brain injury^25^ and pharmacological inhibition of Hdac3’s deacetylase activity reduces neuroinflammation and is protective in models of depression^26^, stroke^23^, and spinal cord injury^24,27^. Hdac inhibition also reduced pro-inflammatory cytokine expression in brains and reversed LPS induced microglial morphology changes^26,28^. However, how Hdac3 regulates gene expression underlying the microglial responses during neuroinflammation is not well understood.

The unique regulation of Hdac3 in microglia supports a critical role for Hdac3 in the regulation of microglial-mediated inflammation and provides a unique opportunity to target Hdac3 clinically to negate the negative impacts of neuroinflammation in brain disease. To investigate the potential neuroprotective mechanisms of Hdac3-inhibition, we investigated epigenetic and gene expression shifts in an *in vitro* microglia model of pro-inflammatory response (LPS).

## Methods

All procedures were approved by the UBC biosafety and ethics committee.

### BV2 Immortalized Microglia Culture

BV2 is a transformed cell line were purchased from ATCC. Cells were cultured in DMEM/F12, 10% HI-FBS, 1x L-Glutamine (ThermoFisher #25030081), 1x Penicillin-Streptomycin. Cells were plated in reduced serum media DMEM/F12 and 2% HI-FBS without antibiotics for each experiment.

### Hdac Inhibitor and LPS Treatments

Hdac Inhibitor drugs RGFP66 (APExBio #A8803) and Suberoylanilide hydroxamic acid (SAHA) (StemCell #73902) were resuspended and stored in DMSO (Cell Signal Technology #12611). BV2 cultures were treated with 15uM RGFP966, 1uM SAHA, or DMSO (vehicle control) for 1 hour.

Lipopolysaccharides (LPS) from *Escherichia coli* (SigmaAldrich #L5418) was diluted to 10ng/ml in distilled H_2_O (vehicle control) and added at concentrations of 0.01-500ng/mL for 1-, 3-, 6-, or 24-hours.

### Reverse Transcription Quantitative Real-Time Polymerase Chain Reaction (RT-qPCR)

BV2 cells were collected in RNA lysis buffer (from Zymo Research Quick-RNA Microprep kit) and RNA extraction performed using Zymo Research Quick-RNA Microprep kit (Zymo Research #R1051). Complementary DNA (cDNA) was synthesized from 200ng RNA using LunaScript^®^ RT-SuperMix kit (New England Biolabs #E3010). RT-qPCR reactions were performed using Luna^®^ Universal qPCR Master Mix (New England Biolabs #M3003). Primers for qPCR reactions are shown in **Table 1**. All primer melt curves were evaluated to verify a single product of the predicted size was produced. Primer efficiency was validated by standard curve. ΔCt values were calculated using the ΔΔCt method. In all experiments, the house keeping gene was tested to verify no significant changes across conditions.

**Table 1.**
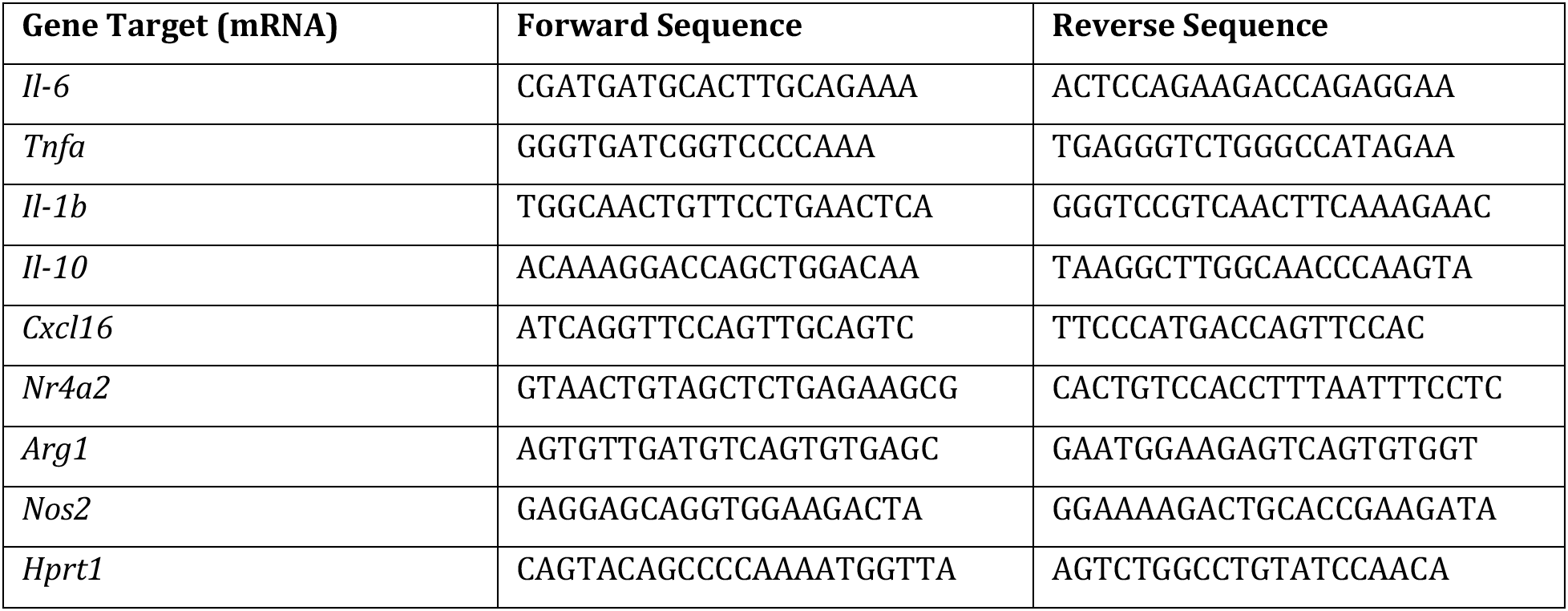
RT-qPCR Primer Sequences.

### Cleavage Under Targets & Release Using Nucleases Assay (CUT&RUN)

CUT & RUN was performed using Cell Signaling Technology CUT & RUN Assay kit (Cell Signaling Technology #86652) according to manufacturer’s protocol. 200,000 Bead-bound cells were incubated with primary antibody overnight at 4°C (1:100 Acetyl-Histone H3 (Lys27) Rabbit mAb (Cell Signaling Technology #8173), 1:50 Acetyl-Histone H3 (Lys9) Rabbit mAb (Cell Signaling Technology #9649), 1:20 negative control Rabbit (DA1E) mAb IgG XP^®^ Isotype Control (Cell Signaling Technology #66362), and 1:50 positive control Tri-Methyl-Histone H3 (Lys4) Rabbit mAb (Cell Signaling Technology #9751)). Bead-cell-antibody samples underwent permeabilization by digitonin buffer, followed by incubation with modified micrococcal nuclease digestion at 4°C for 30 minutes. DNA fragments were released by shaking incubation and isolated using DNA Purification Buffers and Spin Columns (Cell Signaling Technology #14209). Yeast spike-in DNA (5ng) was added to each reaction for normalization using Sample Normalization Primer Set (*Act1)*. Positive control antibody H3K4me3 was tested for successful reaction completion using SimpleChIP^®^ *RPL30* primers. qPCR reactions were performed using Luna^®^ Universal qPCR Master Mix (New England Biolabs #M3003).

### CUT & RUN Primer Design

Primers were designed within the promoter regions (1000bp upstream of transcription start site, TSS) of *Cxcl16, Il-1b, Nos2* and *Arg1* using Primer3Web (Version 4.1.0). Only primers with linear amplification and one product by melt curve analysis were used.

### Input Sample Preparation and Analysis of CUT & RUN qPCR Data

Input control samples were prepared for each treatment condition for whole cell chromatin using micrococcal nuclease (Cell Signaling Technology #10011) digestion to mononucleosomes. qPCR of input samples was run for each qPCR primer set in serial dilutions of 1x, 1:5, 1:25, and 1:125. The Ct values of antibody-isolated CUT & RUN samples were referenced to standard curve (Ct values of input dilutions vs. Log10 (% input)) and calculated as % of input. The % of input for each sample was then normalized by the *Act1* yeast-spike in DNA to account for pipetting error. Normalized values were compared as fold enrichment over the DMSO-H_2_O treatment.

### Phagocytosis Assay Quantified by Flow Cytometry

Phagocytic activity of BV2 microglia was detected using engulfment of pHrodo Red *E.coli* BioParticles ™ Conjugate for Phagocytosis (ThermoFisher Scientific #P35361). Cells were treated in last hour of incubation with 1:500 dilution of pHrodo red *e.coli* BioParticles ™. Cells were washed with FACS buffer and resuspended in 1% PFA for overnight 4°C fixation. The next day cells were washed twice in FACS buffer and run on the CytoFLEX Flow Cytometer. Flow cytometry data was analyzed using gating for cell size, granularity, singlet cell population, and phycoerythrin (PE) red-channel signal to detect cells with bead engulfment. FlowJo was used to assess the percent of phagocytic positive cells gated on the no stain in the PE channel and the median fluorescent intensity (MFI) of the positive population.

### Protein Quantified by Flow Cytometry

Global protein concentration of the BV2 cells post treatment was assessed via flow cytometry using intracellular protein staining using the True-Nuclear Transcription Factor Buffer Set (Biolegend # 424401). Cells were incubated with primary antibodies for 30 minutes - 1:100 Acetyl-Histone H3 (Lys27) Rabbit mAb (Cell Signaling Technology #8173) or 1:250 Acetyl-Histone H3 (Lys9) Rabbit mAb (Cell Signaling Technology #9649). Antibodies were detected with 1:500 AlexaFluor^®^ 568 Donkey Anti-Rabbit (Invitrogen #A10042) incubated with cells for 30 minutes. Cells were run on the CytoFLEX Flow cytometer. FlowJo was used to gate the cells for cell size (FSC A vs SSC A), singlets (FSC-H vs FSC-W), and then for positive signal in the 585 channel to detect antibody fluorescence. MFI for the 585 positive population was used as a measure for protein level. MFIs were normalized to the control condition to determine fold change and compared across conditions.

### Griess Reagent Assay

The Griess Reagent kit (ThermoFisher #G7921) was used to quantify nitrite concentrations in media released by BV2 microglia as described by the manufacturer. A standard curve was prepared from nitrite-containing samples and used to determine sample concentrations.

### Statistical Analysis

In instances of one treatment, ordinary one-way ANOVAs were run comparing the mean of each treatment to the control. Dunnett’s post hoc comparisons were run for individual treatment comparisons. Residuals were tested for normality using Shapiro-Wilk test. In instances of two treatments (Hdac inhibitor and LPS treatment) a two-way ANOVA was run to fit a full effect model (Hdac inhibitor, LPS treatment and the interaction). Tukey’s or Sidak post hoc comparisons were run to compare individual conditions. Residuals were tested for normality using Shapiro-Wilk test. All measures passed normality testing.

## Results

### BV2 Microglial Cells Response to LPS Treatment

Previous work has assessed the feasibility of BV2 microglia as a robust and model of microglial responses^29,30^. We initially performed a dose curve experiment to assess the gene expression and histone acetylation responses in BV2s to LPS. The expression of interleukin-6 (*Il-6)* (F(3,8)=122.0, p<0.0001), tumor necrosis factor alpha (*Tnfa*) (F(3,8)=32.95, p<0.0001), and interleukin 1 beta (*Il-1b*) (F(3,8)=58.18, p<0.0001), were all significantly increased in expression at an LPS range of 10 – 500ng/mL^31^ for a 3-hour duration (**Figure 1A**).

**Figure 1.**
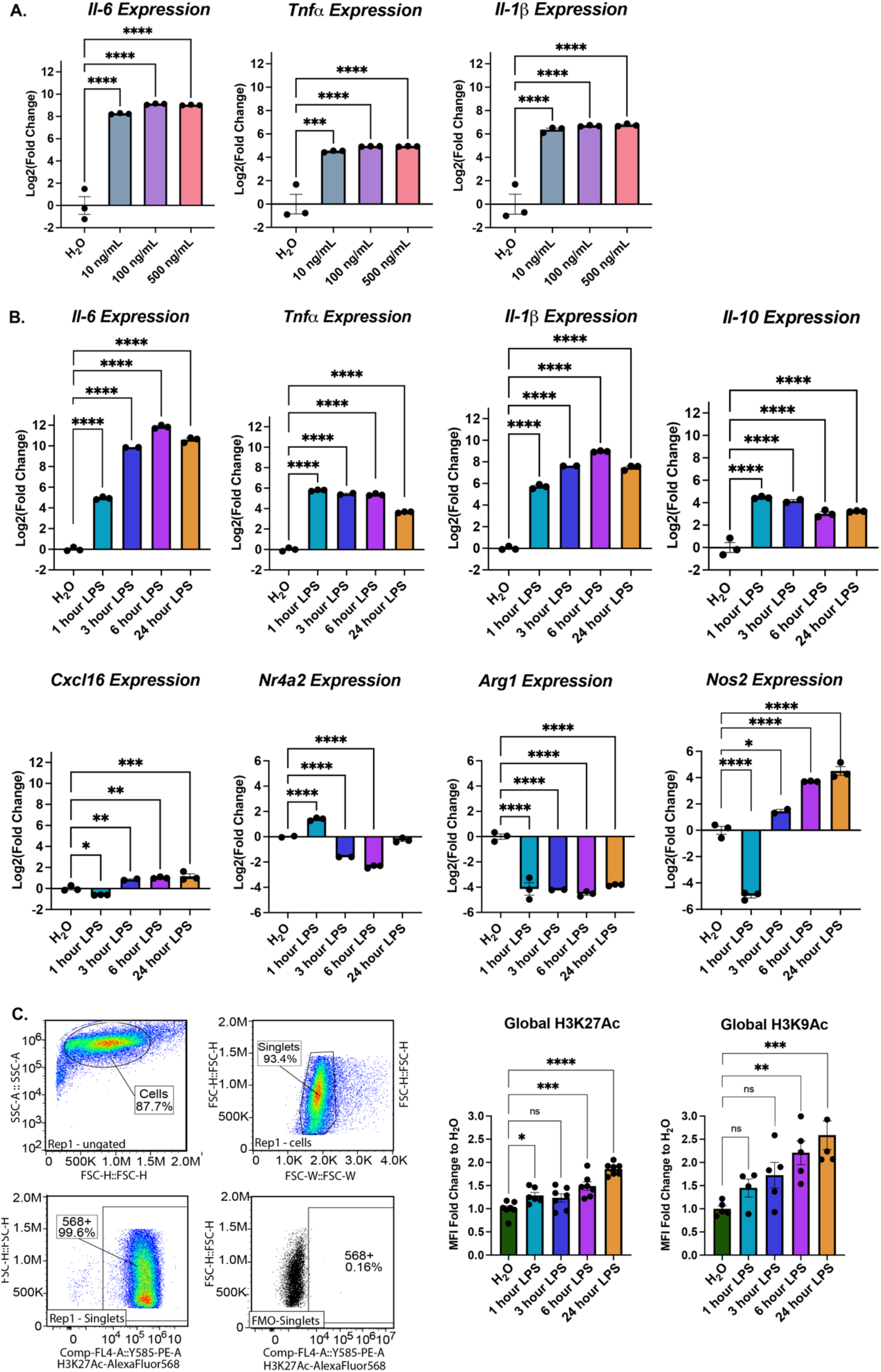
BV2 Immortalized microglia robustly regulate gene expression and histone acetylation to LPS treatment. (A) RT-qPCR assessment of gene expression of *Il-6, Tnfa*, and *Il-1b* in BV2 microglia treated with different LPS doses 10, 100, or 500ng/mL for 3 hours. Shown as bar graph of Log2(Fold Change) + SEM Dunnett’s post hoc significances denoted (*** p<0.0002, **** p<0.0001). (B) RT-qPCR assessment of gene expression of *Il-6, Tnfa, Il-1b, Il-10, Cxcl16, Nr4a2, Arg1*, and *Nos2* of BV2 microglia treated with 10ng/mL LPS for 1, 3, 6, or 24 hours. Shown as bar graph of Log2(Fold Change) + SEM Dunnett’s post hoc significances denoted (*p<0.03, ** p<0.002, *** p<0.0002, **** p<0.0001). (C) Global histone modifications by flow cytometry. Events are gated for cell size on side scatter (SSC) Area vs Forward Scatter (FSC) height and singlets on FSC-height vs FSC-width. The cells that are positive for the histone mark signal are gated based on an FMO in the same channel. The median fluorescence intensity of the positive population is exported for downstream analysis. Median fluorescent intensity (MFI) for global levels of H3K27ac levels as measured by intracellular flow cytometry. Fold Change + SEM. Dunnett’s post hoc significances denoted (*p<0.05, ** p<0.007, *** p<0.0005, **** p<0.0001). n=2-7 replicates per condition from at least two independent sets of cultures.

Responsiveness to 10ng/mL LPS was then tested over a time course of 1, 3, 6, and 24 hours (**Figure 1B**) for gene expression of pro-inflammatory cytokines *Il-6, Tnfa, Il-1b*, anti-inflammatory cytokine *Il-10*, chemokine (C-X-C motif) ligand 16 (*Cxcl16*), nuclear receptor subfamily 4 group A member 2 (*Nr4a2*), arginase 1 (*Arg1*), and nitric oxide synthase 2 (*Nos2*). One-way ANOVAs revealed a significant main effect of LPS duration for *Il-6* (F(4,9)=2293, p<0.0001), *Tnfa* (F(4,9)=1410, p<0.0001), *Il-1b* (F(4,9)=1379, p<0.0001), and *Il-10* (F(4,9)=58.51, p<0.0001), *Cxcl16* (F(4,9)=32.45, p<0.0001), *Nr4a2* (F(4,8)=405.4, p<0.0001), *Arg1* (F(4,9)=52.09, p<0.0001), and *Nos2* (F(4,9)=269.2, p<0.0001). Dunnett’s post hoc comparisons were run for each LPS duration compared to H2O control and revealed the expected significant increase in cytokine and chemokine expression by 1hr of treatment that largely maintained to 24hrs. *Nr4a2*, a known Hdac3 target gene in neurons, showed a transient increase at 1hr followed by a significant repression at subsequent time points. *Arg1* and *Nos2*, two enzymes that regulate nitric oxide (NO) production, showed significant repression at 1hr of LPS. *Arg1* remained below baseline levels while *Nos2* increased above baseline by 6 and 24 hours of treatment.

To examine impact of LPS treatment on histone acetylation, we measured global protein levels of H3K27ac and H3K9ac by intracellular flow cytometry (**Figure 1 C**). LPS treatment significantly increased both H3K27ac (F(4,29)=19.61, p<0.0001) and H3K9ac (F(4,19)=6.892, p=0.0013) H3K27ac was significantly increased by 1 hour and H3K9ac after 6 hours of LPS.

### Hdac Inhibition Modulates BV2 Microglial Gene Expression

To test the role for Hdac3 in regulating LPS mediated gene expression, we utilized the Hdac3 selective small molecule inhibitor RGFP966 that inhibits the enzymatic activity of Hdac3 (IC_50_=80nM) with >200-fold selectivity over other Hdacs^21^. As a comparison, we also used the clinically approved pan-Hdac inhibitor SAHA that inhibits all Class I and II HDACs (HDAC 1-10)^32^. Hdac inhibitors were applied to BV2 cultures for 1 hour prior to treatment with either LPS (10ng/ml) or vehicle (water) for 1 or 3 hours (**Figure 2A**). We then examined the mRNA expression of *Il-1b, Tnfa, Il-10, Cxcl16, Nr4a2, Arg1*, and *Nos2*.

**Figure 2.**
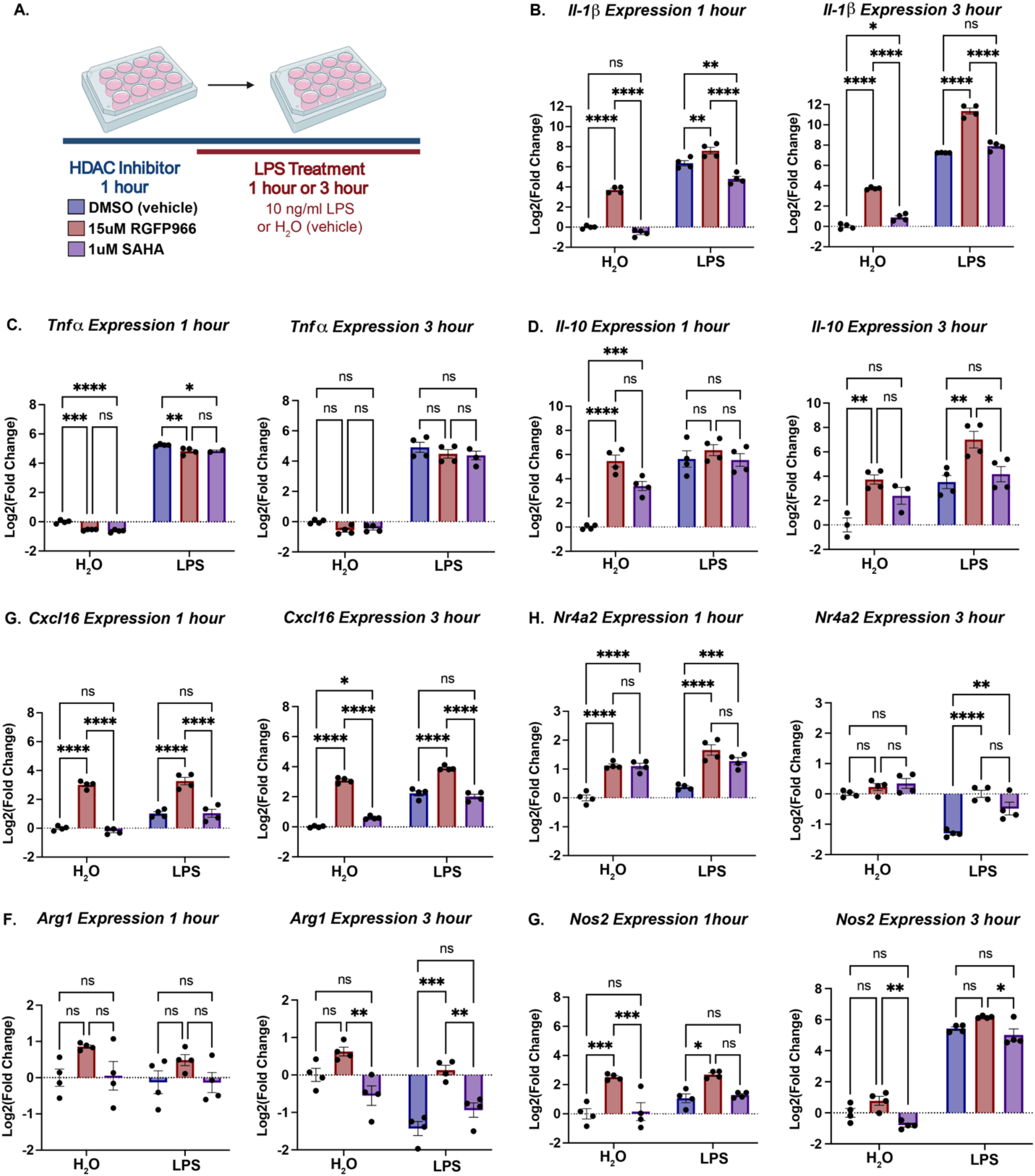
Hdac inhibition modulates LPS regulated gene expression. (A) Experimental design for testing the role of Hdacs in modulating LPS regulated gene expression in BV2 cultures. For all subsequent panels DMSO control is shown in blue, RGFP966 shown in red and SAHA in purple. (B) *Il-1b* expression is enhanced by Hdac3 inhibition at baseline and in response to LPS at 1 and 3 hours of treatment. (C) Tnfa expression is slightly repressed by Hdac inhibition at 1 hour and unmodulated at 3 hours of LPS treatment. (D) *Il-10* expression is enhanced by Hdac3 inhibition at baseline and in response to LPS at 3 hours of treatment. (E) *Cxcl16* expression is enhanced by Hdac3 inhibition at baseline and in response to LPS at 1 and 3 hours of treatment. (F) *Nr4a2* expression is enhanced by both RGFP966 and SAHA at baseline and in response to LPS at 1 hour. Both RGFP966 and SAHA prevent the normal, LPS induced repression of *Nr4a2* expression at 3 hours of LPS. (G) *Arg1* expression is not above baseline in any condition at 1 hour of LPS treatment. At 3 hours, RGFP966, but not SAHA, blocks LPS induced repression of *Nr4a2*. (H) At 1 hour LPS, *Nos2* expression is increased by Hdac3 inhibition, but not SAHA, under baseline conditions and in response to LPS. At 3 hours expression largely matches DMSO treated controls. Each panel is Log2 fold change relative to DMSO control treated with water and error bars are +/- SEM. In all panels DMSO water versus DMSO LPS significance is not shown, but reaches statistical threshold (*p<0.05) for all genes at 1 hour of LPS except *Nr4a2, Arg1*, and *Nos2* and was significant for all genes at 3 hours LPS. n=3-4 per condition in at least 3 independent replication experiments. *p<0.05, **p<.01, ***p<0.001, ****p<0.0001.

A two-way ANOVA for *Il-1b* at 1 hour of LPS treatment revealed a significant main effect of LPS (F(1,18)=834.6, p<0.001), of Hdac inhibition (F(2,18)=135.1, p<0.0001) and an interaction F(2,18)=15.55, p=0.0001). At 3 hour of LPS treatment, *Il-1b* showed a significant main effect of LPS (F(1,18)=2638, p<0.001), of Hdac inhibition (F(2,18)=285.7, p<0.0001) and but no interaction F(2,18)=1.461, p=0.2581). Tukey’s multiple comparisons revealed a significant increase in *Il-1b* with RGFP966 above vehicle both at baseline and in response to LPS at 1 and 3 hours. This enhancement in expression was not observed with SAHA treatment at 1 hour and a slight increase in baseline at 3 hours, suggesting the increased expression is largely driving by inhibition of Hdac3.

We next investigated the expression of *Tnfa* and found at 1 hour LPS a significant main effect of LPS (F(1,16)=9439, p<0.0001), Hdac inhibition (F(2,16)=39.72, p<0.001) but no interaction (F(2,16)=1.339, p=0.2899). At three hours there was a main effect of LPS (F(1,18)=728.8, p<0.0001) but no effect of Hdac inhibition (F(2,18)=3.308, p=0.0598) nor interaction (F(2,18)=0.1194, p=0.8881). Tukey’s posthoc comparisons at 1 hour LPS showed a small but significant decrease in *Tnfa* expression at baseline and with LPS for both RGFP966 and SAHA. All comparisons were no longer significant by 3 hours, indicating that Hdacs are not negative regulators of *Tnfa* expression.

We also examined expression of the anti-inflammatory cytokine *Il-10* and similar to *Il-1b*, we observed largely RGFP966 specific enhancements in gene expression. At 1 hour of LPS treatment there was a significant main effect of LPS (F(1,18)=56.88, p<0.0001), Hdac inhibition (F(2,18)=21.69, p<0.0001) and a significant interaction (F(2,18)=13.64, p=0.0002). At 3 hours of LPS treatment there was a significant main effect of LPS (F(1,16)=34.72, p<0.0001), Hdac inhibition (F(2,16)=19.24, p<0.0001) and but no interaction (F(2,16)=1.228, p=0.3191). Tukey’s posthocs revealed a significant increase in *Il-10* levels at baseline with both RGFP966 and SAHA, but no modulation of LPS induction. In contrast, at 3 hours of LPS there was a significant increase in baseline *Il-10* expression and enhanced response to LPS only with RGFP966. These findings indicate that both pro and anti-inflammatory cytokines are modulated similarly by Hdac3 inhibition.

To further explore the impact on additional gene targets, we examined the chemokine *Cxcl16*. Similar to *Il-1b* and *Il-10*, for *Cxcl16* expression 1 hour of LPS treatment revealed a significant main effect of LPS (F(1,18)=33.53, p<0.0001), Hdac inhibition (F(2,18)=151.7, p<0.0001) and a significant interaction (F(2,18)=4.242, p=0.0310). At 3 hours of LPS treatment, there was also a significant effect of LPS (F(1,18)=286.0, p<0.0001), Hdac inhibition (F(2,18)=302.0, p<0.0001) and an interaction (F(2,18)=21.56, p<0.0001). Tukey’s posthocs revealed a significant increase in *Cxcl16* expression both at baseline at both time points with Hdac3 inhibition but and minimal impact with SAHA. In response to LPS, only the Hdac3 inhibitor enhanced gene expression beyond levels observed in the DMSO controls, suggesting that Hdac3 may be the predominant regulator of *Cxcl16*.

In contrast, *Nr4a2* expression was modulated by both RGFP966 and SAHA. At 1 hour of LPS, there was a significant main effect of LPS (F(1,18)=16.23, p=0.0008) and Hdac inhibition (F(2,18)=68.63, p<0.0001) but no interaction (F(2,18)=1.404, p=0.2713). At 3 hours of LPS, there was a significant main effect of LPS (F(1,18)=54.16, p<0.0001) and Hdac inhibition (F(2,18)=18.69, p<0.0001) and an interaction (F(2,18)=8.534, p=0.0025). Tukey’s posthocs revealed a significant increase in baseline gene expression with both RGFP966 and SAHA at 1 hour of LPS but not 3 hours. In response to LPS at 1 hour there was a significant increase above DMSO-LPS for both Hdac inhibitors. At 3 hours of LPS, the DMSO control showed decreased gene expression relative to baseline, but this repression failed to occur in both Hdac inhibitor treated conditions. Together, this indicates that Nr4a2 is potentially regulated by multiple Hdacs in a bi-directional manner with enhanced baseline expression and impaired LPS mediated repression.

To examine another gene with LPS induced repression, we measured *Arg1* gene expression. At 1 hour of LPS there was not a significant main effect of LPS (F(1,18)=1.134, p=0.3010), a significant main effect of Hdac inhibition (F(2,18)=5.057, p=0.0181) and no interaction (F(2,18)=0.1193, p=0.8882). At 3 hours of LPS, there was a significant effect of LPS (F(1,18)=26.81, p<0.0001) and of Hdac inhibition (F(2,18)=24.62, p<0.0001) and a significant interaction (F(2,18)=4.994, p=0.0188). Tukey’s posthocs revealed no significant differences at 1 hour of LPS. At 3 hours, there was no difference at baseline for either Hdac inhibitor compared to vehicle control, but a trend for a decrease with SAHA. In response to 3 hours of LPS, there was a significant repression in the DMSO control and SAHA treated samples, but not in the RGFP966 sample treated with LPS, similar to the failure in gene repression observed for *Nr4a2*.

As Arg1 acts in opposition to iNos (*Nos2*) in the generation of nitric oxide, we also examined impacts on *Nos2* expression. At 1 hour of LPS treatment there was a significant main effect of LPS (F(1,18)=8.917, p=0.0079) and Hdac inhibition (F(2,18)=25.00, p<0.0001), but no significant interaction (F(2,18)=1.364, p=0.2809). Similarly, at three hours of LPS, there was a significant main effect of LPS (F(1,18)=801.1, p<0.0001) and of Hdac inhibition (F(2,18)=16.18, p<0.0001), but no interaction (F(2,18)=0.4633, p=0.6365). Tukey’s posthoc comparisons revealed a significant increase in *Nos2* expression at baseline and in response to LPS at 1 hour, an effect not observed with SAHA. At three hours, there was no difference between either Hdac inhibitor treated condition and the respective DMSO controls, indicating the impacts on *Nos2* expression are short lived.

### Hdac Inhibition Enhances Histone Acetylation

The Hdac inhibitor impacts on BV2 microglial gene expression both at baseline and in response to LPS, support a model in which increased histone acetylation is permissive for LPS regulated gene expression. Consequent increases in histone acetylation upon Hdac inhibition would then be consistent with the observed pattern of released gene repression at baseline, enhanced LPS induced gene expression and prevention of LPS induced repression of gene expression. To test this prediction, we first examined global histone acetylation changes in response to Hdac inhibition and LPS treatment by flow cytometry. Global levels of H3K9ac showed a significant main effect of Hdac inhibitor (F(2,45)=104.18, p<0.0001), LPS treatment (F(1,45)=11.50, p=0.0015), but no interaction (F(2,45)=1.304, p=0.2814). H3K27ac also showed a robust increase upon Hdac inhibition with a significant main effect of Hdac inhibitor (F(2,44)=77.77, p<0.0001), LPS treatment (F(1,44)=6.841, p=0.0122), and no significant interaction (F(2,44)=2.999, p=0.0601). Tukey’s corrected posthoc comparisons revealed that both Hdac inhibitors significantly increased H3K9ac and H3K27ac at baseline and upon LPS treatment (**Figure 3**).

**Figure 3.**
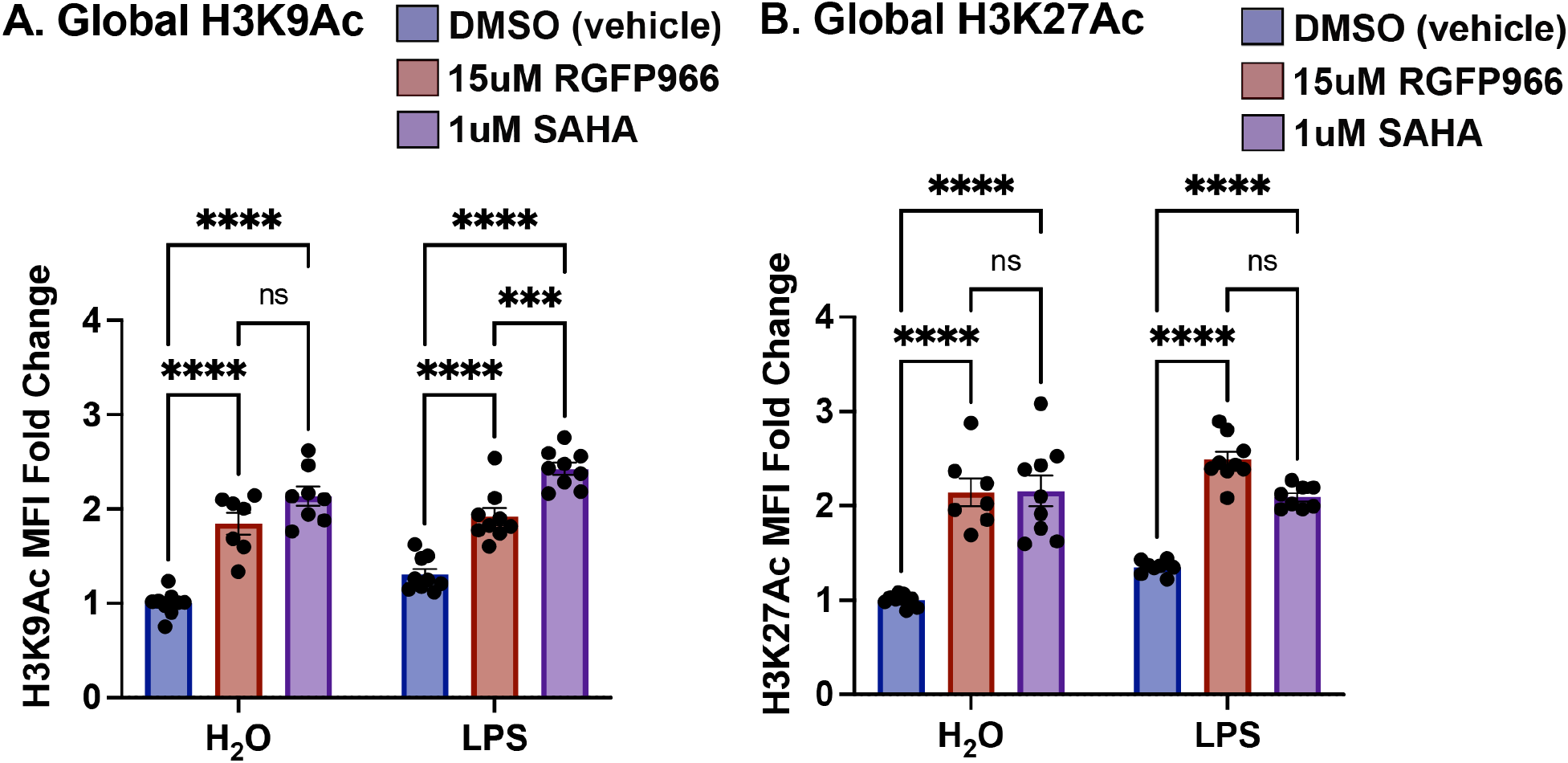
Global changes in histone acetylation upon Hdac inhibition. BV2 microglial cells were treated with DMSO, RGFP966 or SAHA for 1 hour and then either water or LPS was added for three hours. Cells were then harvested and intracellular staining was performed for H3K9ac or H3K27ac. (A) Global levels of H3K9ac were significantly increased with both RGFP966 and SAHA at baseline and upon LPS treatment. (B) H3K27ac levels were increased with both RGFP966 and SAHA at baseline and upon LPS treatment. The magnitude of increase was similar with the two Hdac inhibitors. Tukey’s corrected posthocs *p<0.05, **p<0.005, ***p<0.001, ****p<0.0001. MFI: median fluorescence intensity. Fold change is relative to DMSO treated water samples. n=5-6 per treatment in 3 independent replication experiments.

To examine the link between gene expression and histone acetylation we examined H3K27ac and H3K9ac by CUT&RUN qPCR at the promoters of select genes with expression regulated by Hdac3 inhibition (**Figure 4A**). The positive control H3K4me3 antibody produced significant enrichment over non-immune IgG (t(6)=2.511, p=0.0458) indicating the CUT&RUN procedure was working as expected. We examined H3K27ac and H3K9ac over the Cxcl16 promoter (**Figure 4B**). There was a significant main effect of Hdac inhibition (F(1,8)=18.76, p=0.0025) but not for LPS treatment (F(1,8)=0.3512, p=0.5698) nor for the interaction (F(1,8)=0.0003, p=0.9576). Sidak corrected posthoc comparisons between RGFP966 and DMSO treated samples revealed a significant increase in H3K27ac promoter signal with Hdac3 inhibition at both baseline and in response to LPS. For H3K9ac over the Cxcl16 promoter, there was a significant main effect of Hdac inhibition (F(1,8)=33.86, p=0.0004) and for LPS treatment (F(1,8)=6.101, p=0.0387), but not for the interaction (F(1,8)=0.2133, p=0.6565). Sidak corrected posthoc comparisons revealed a significant increase in H3K9ac promoter signal with Hdac3 inhibition at both baseline and in response to LPS.

**Figure 4.**
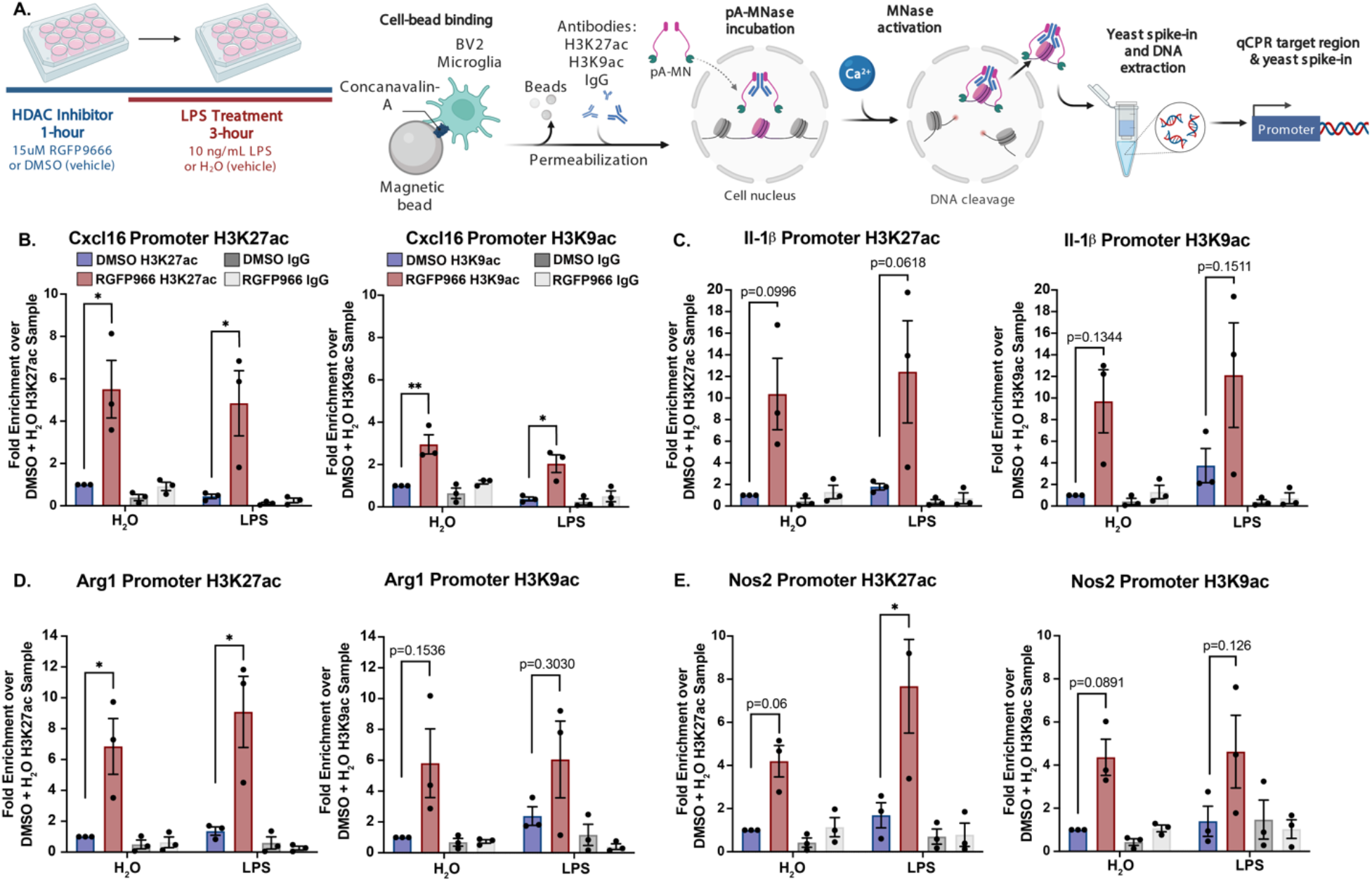
Histone acetylation changes at Hdac3 modulated genes. (A) Experimental design for CUT&RUN qPCR analysis of H3K27ac and H3K9ac over promoters of key Hdac regulated target genes. All samples are normalized to yeast spike in and expressed as a percentage of input sample based on standard curve. Signal then calculated as fold enrichment relative to the H3K27ac or H3K9ac antibody levels in the DMSO and water treated control. DMSO and RGFP966 non-immune IgG samples were included for all experiments (B) H3K27ac and H3K9ac signal is significantly increased over the Cxcl16 promoter with RGFP966 treatment. (B) Trends for increased H3K27ac and H3K9ac signal with RGFP966 over the Il1-b promoter. (C) H3K27ac signal is significantly increased over the Arg1 promoter with RGFP966 treatment. H3K9ac shows similar trends but did not reach significance. (D) H3K27ac signal is significantly increased over the Nos2 promoter with RGFP966 and LPS treatment. H3K9ac shows similar trends but did not reach significance. All plots are fold enrichment with +/- SEM. n=3 per condition in independent replication experiments. *p<0.05.

For the Il1-b promoter H3K27ac levels, we found a significant main effect of Hdac inhibition (F(1,8)=11.99, p=0.0085), but no effect of LPS (F(1,8)=0.2425, p-0.6356) nor interaction (F(1,8)=0.048, p=0.8314). H3K9ac levels showed a similar pattern over the Il1b-promoter with a significant main effect of Hdac inhibition (F(1,8)=8.440, p=0.0187), no effect of LPS (F(1,8)=0.773, p=0.4047) nor interaction (F(1,8)=0.003, p=0.9571). Sidak corrected posthocs revealed trends towards enhanced H3K27ac signal over the promoter at baseline and with LPS (**Figure 4C**).

At the Arg1 promoter, H3K27ac levels showed a significant main effect of Hdac inhibition (F(1,8)=21.25, p=0.0017), no effect of LPS (F(1,8)=0.7804, p=0.4028) and no interaction (F(1,8)=0.4074, p=0.5411). For H3K9ac, there was a significant main effect of Hdac inhibition (F(1,8)=6.237, p=0.0371), no effect of LPS (F(1,8)=0.2290,p=0.6451) and no interaction (F(1,8)=0.1136, p=0.7447). Sidak corrected posthocs revealed a significant increase in H3K27ac signal with RGFP966 over DMSO at both baseline and in response to LPS. Similar trends in increase were also observed for H3K9ac, but did not reach significance (**Figure 4D**). At the Nos2 promoter, H3K27ac levels showed a significant main effect of Hdac inhibition (F(1,8)=15.10, p=0.0046), no effect of LPS (F(1,8)=3.112, p=0.1157) and no interaction (F(1,8)=1.392, p=0.2719). For H3K9ac, there was a significant main effect of Hdac inhibition (F(1,8)=10.75, p=0.0112), no effect of LPS (F(1,8)=0.1060,p=0.7531) and no interaction (F(1,8)=0.0043, p=0.9493). Sidak corrected posthocs revealed a significant increase in H3K27ac with RGFP966 upon LPS treatment and a trend at baseline. There were trends for RGFP966 induced increase in H3K9ac at baseline and with LPS, but they did not reach statistical significance (**Figure 4E**).

### Hdac Inhibition Enhances Microglial Phagocytosis and Impairs NO Release

To examine how Hdac inhibition may impact microglia function, we examined phagocytosis of pH-rodo E. coli labelled beads. These beads specifically fluoresce when in the low pH environment of the phagolysosome and can then be quantified by flow cytometry. We quantified the impact of Hdac inhibition and LPS treatment on both the percentage of microglia that phagocytose beads and median fluorescence intensity (MFI) of the engulfed beads, a proxy for the number of beads phagocytosed (**Figure 5A**). At three hours of LPS treatment there was no significant impact of LPS (F(1,36)=0.0796, p=0.7795) or Hdac inhibition (F(2,36)=3.119, p=0.563) or an interaction F(2,36)=0.1551, p=0.8569) for the percentage of bead positive microglia. For the MFI there was no significant effect of LPS (F(1,36)=3.521, p=0.0687) but there was a significant effect of Hdac inhibition (F(2,36)=7.990, p=0.0013), and no interaction F(2,36)=0.5893, p=0.5600) (**Figure 5B**). At 24 hours of LPS treatment, there were more significant impacts on microglial phagocytosis. The percentage of positive microglia showed a significant effect of LPS F(2,36)=21.64, p<0.0001) and Hdac inhibition (F(1,36)=14.28, p=0.0006) and a significant interaction (F(2,36)=19.21, p<0.0001). For the MFI at 24 hours there was no effect of LPS (F(1,36)=0.0004, p=0.9851), but there was a significant effect of Hdac inhibition F(2,36)=16.31, p<0.0001) and a significant interaction F(2,36)=13.92, p<0.0001) (**Figure 5C**). Tukey’s corrected posthoc comparisons revealed a significant increase in phagocytosis positive microglia with both RGFP966 and SAHA treatment at baseline. There was no difference between Hdac inhibitor treatments with LPS, indicating that Hdac inhibition may drive maximal phagocytosis even without immune stimulation. At 24 hours of LPS treatment, Hdac inhibition produced a significant enhancement in the amount of phagocytosis of individual microglia at baseline but not in response to LPS.

**Figure 5.**
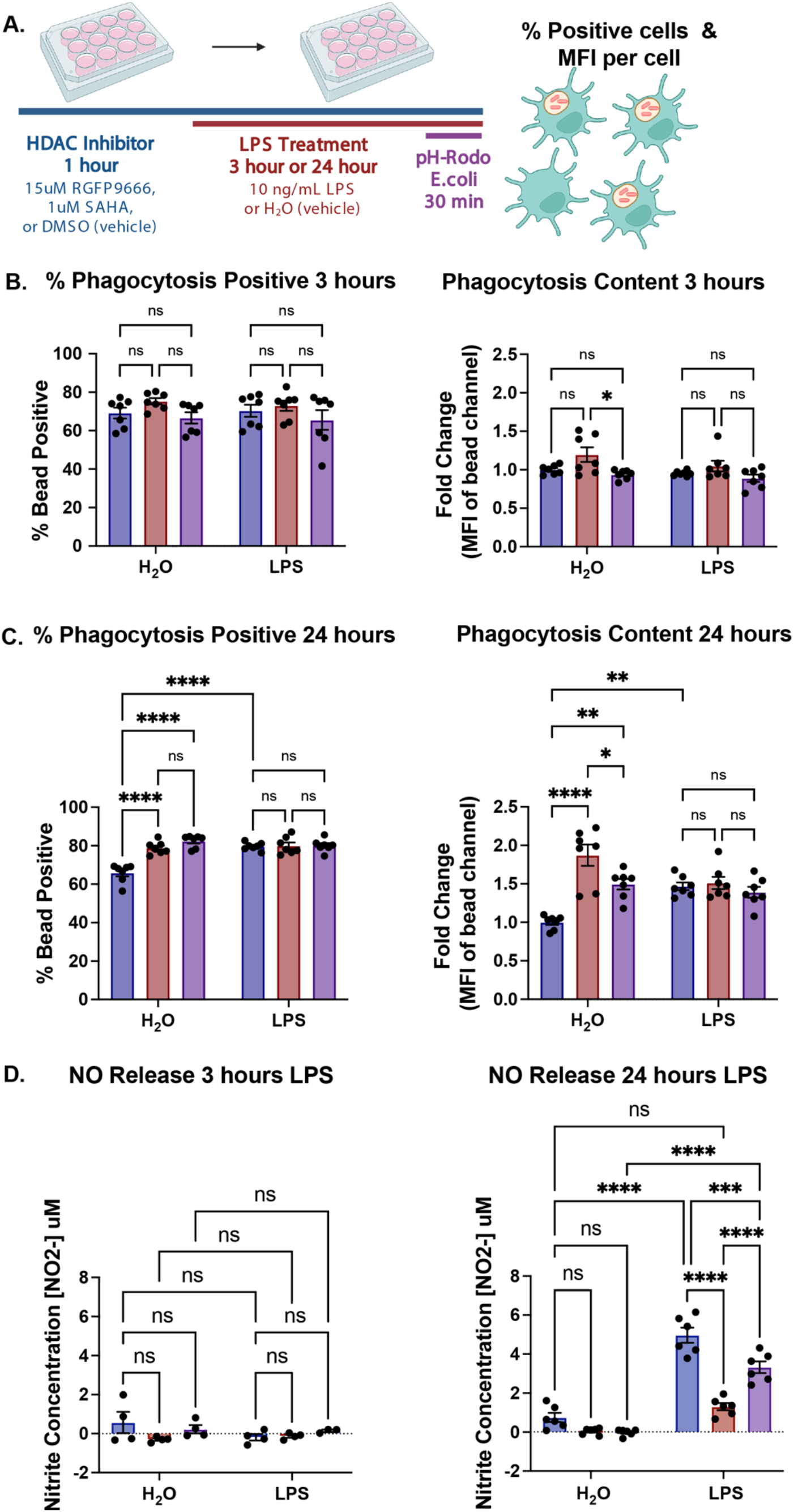
Hdac inhibition enhances microglial phagocytosis and suppresses nitric oxide release. (A) Experimental design for phagocytosis assay. Both the percent of positive microglia and the MFI per microglia are quantified by flow cytometry (B) Hdac inhibition shows minimal impacts on phagocytosis of pH-rodo *E. coli* tagged beads at 3 hours of LPS treatment. (C) Hdac inhibition enhances phagocytosis at baseline after 24 hours, bringing the percent of positive microglia to LPS levels. (D) Hdac inhibition enhances phagocytosis at baseline after 24 hours, bringing it above LPS induced levels in the case of RGFP966. (E) Minimal release of NO after 3 hours of LPS. (E) At 24 hours of LPS treatment, DMSO treated samples show a significant increase in NO release. This effect was blunted with SAHA and completely repressed to baseline levels with RGFP966. Mean +/- SEM. n=4-7 from 2 or 3 independent experiments. *p<0.05, **p<.01, ***p<0.001, ****p<0.0001.

To further examine microglia function we measured release of NO into the media both at 3 hours and 24 hours of LPS treatment. At 3 hours, there were no significant differences between conditions and overall levels of NO were low. There was no effect of LPS (F(1,17)=1.033, p=0.3237), Hdac inhibition (F(2,17)=1.565, p=0.2337) nor interaction (F(2,17)=1.691, p=0.2140). At 24 hours there was a significant effect of LPS (F(1,30)=226.9, p<0.0001) of Hdac inhibition (F(2,30)=41.95, p<0.0001) and a significant interaction (F(2,30)=20.23, p<0.0001). Tukey’s corrected posthocs revealed a significant increase in NO release upon LPS treatment in the DMSO condition. This increase was blunted with SAHA treatment and reduced to baseline levels with RGFP966 (**Figure 5D**).

## Discussion

Pharmacological inhibition of Hdac3’s deacetylase activity reduces neuroinflammation and is protective in models of depression^26^, stroke^23^, and spinal cord injury^24,27^. Conditional deletion of Hdac3 in microglia shifted microglial responses to a traumatic brain injury towards a more inflammation resolving phenotype and improved functional recovery^25^. Together this suggests a pro-inflammatory role for Hdac3 in regulating microglial function and that suppression of Hdac3 is beneficial for combating neuroinflammation. Our findings indicate that the beneficial effects of inhibiting Hdac3 may be due to several functions of Hdac3 in microglia. We found that Hdac3 inhibition robustly increased histone acetylation and gene expression of numerous cytokines (*Il1- b, Il-10*), chemokines (*Cxcl16*) and LPS inducible regulators (*Nr4a2, Arg1, Nos2*) both at baseline and in response to LPS. Hdac3 inhibition also enhanced phagocytosis while simultaneously blunting NO release. While classically considered a pro-inflammatory response, enhanced phagocytosis following injury or acutely during disease is often beneficial for clearing dead or dying cells in the brain. Augmenting this microglial response may ultimately facilitate brain recovery after damage. RGFP966 suppression of NO release from microglia may further promote resolution of inflammation by damping downstream NO induced pro-inflammatory signals.

Our gene expression findings indicate that RGFP966 may regulate NO release through control of expression of *Arg1* and *Nos2* (iNos). The enzymatic processes of Arg1 and iNos compete for the substrate L-arginine with opposing cellular phenotypes ^33^. Arg1 hydrolyzes L-arginine to produce urea and L-ornithine, which removes nitrogen from amino acid metabolism via the urea cycle and promotes cell proliferation^34^. The substrate L-arginine is also used by iNos in the production of L-citrulline and NO. We found a decrease in *Arg1* expression with LPS treatment across timepoints. This would effectively decrease competition for L-arginine, allowing iNos to increase production of NO, as we observed in DMSO treated cells upon LPS. RGFP966 effectively prevented the decrease in *Arg1* expression and had only marginal impacts on *Nos2*, potentially shifting the microglial activation state away from NO production (**Figure 6**). This would be consistent with the overall protective effect of RGFP966 in the context of stroke^23^ and spinal cord injury^24,27^.

**Figure 6.**
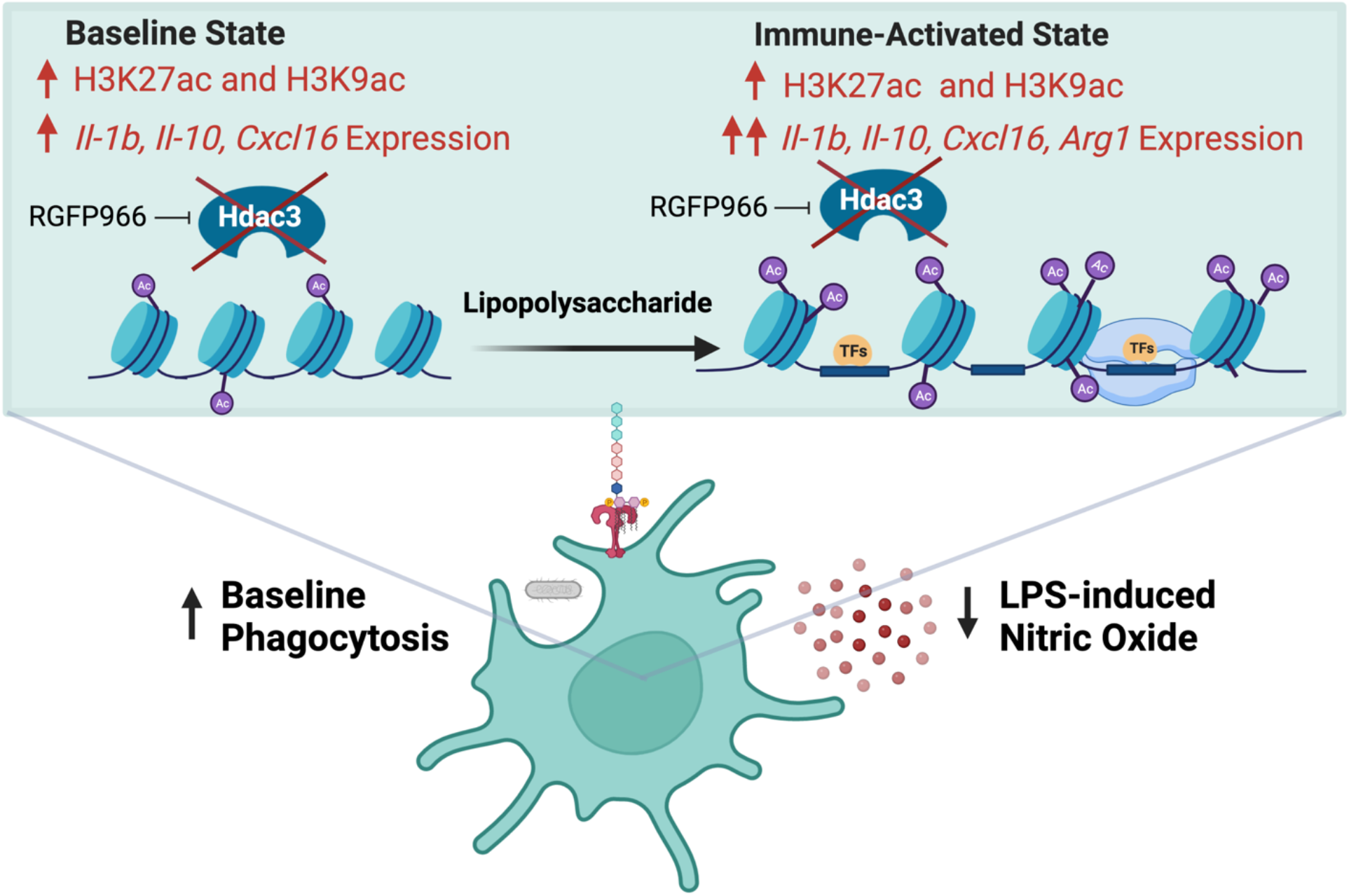
Model for Hdac3 regulation of microglial LPS mediated gene expression and function. Hdac3 represses baseline gene expression through deacetylation of H3K27ac and H3K9ac marked histones. Upon LPS stimulation Hdac3 releases, allowing for increased histone acetylation and immune activation. In the presence of RGFP966, Hdac3’s deacetylase activity is blocked allowing for increased histone acetylation at baseline and an aberrant increase in baseline gene expression. Upon LPS, inhibition of Hdac3 results in hyper-induction of Hdac3 target genes and a lack of suppression of genes normally repressed by LPS treatment. These gene expression shifts ultimately culminate in increased phagocytosis and repressed NO release, driving microglia towards a phenotype that promotes resolution of inflammation.

In peripheral macrophages loss of Hdac3 produces complex impacts on gene expression. Loss of Hdac3 in lung macrophages or cultured bone marrow derived macrophages (BMDM) increases expression of genes that promote wound-healing^38^. Similar to our findings in microglia, Hdac3 inhibition significantly increased baseline expression of genes that were normally downregulated by LPS stimulation in BMDM^42^. Paradoxically, macrophage Hdac3 also promotes activation of inflammatory gene expression^39,43^ through a non-canonical activating role via recruitment of activating transcription factor ATF2. The activating role does not require Hdac3’s deacetylase activity, and consequently genetic deletion of Hdac3 results in loss of both the canonical transcriptional repression and the non-canonical transcriptional activating roles. Our findings in microglia only directly tested the enzymatic role of Hdac3 in regulating gene expression and future work with conditional deletions of Hdac3 will be required to identify if microglial Hdac3 also has a non-canonical activating function.

One of the limitations of our study is that all experiments were performed in the BV2 microglial immortalized cell line. While *in vitro* studies provide a number of advantages for high throughput and controlled testing of gene expression mechanisms, *in vitro* regulation does not always recapitulate *in vivo* microglial regulation^3,50^. Henn et. al (2009)^29^ found that the majority of genes induced in BV2 cells by LPS treatment were also induced in primary microglia (90%) and freshly isolated hippocampal microglia (50%), although BV2 microglia gene expression changes were less pronounced than primary microglia. Given similar functional findings using RGFP966 *in vivo* and recent findings showing altered microglial responses in Hdac3 microglial conditional knockout mice^23–27^, we believe our *in vitro* model captures fundamental gene regulation mechanisms and demonstrates how Hdac3 regulation of microglial gene expression leads to *in vivo* improvements in models with brain inflammation.

## Conclusion

Together our findings demonstrate an important role for Hdac3 as a negative regulator of the microglial gene expression response to LPS. Our epigenetic profiling indicates Hdac3 suppresses microglial gene expression through deacetylation of histone targets at the promoters of both classically pro- and anti-inflammatory genes. Inhibition of Hdac3 shifts the microglial LPS response towards resolution of inflammation through enhanced phagocytosis and reduced NO release. Our findings support a model in which Hdac3 inhibition driven shifts in microglial gene expression and function ultimately conveys neuro-protection in brain disease.

**Table 2.**
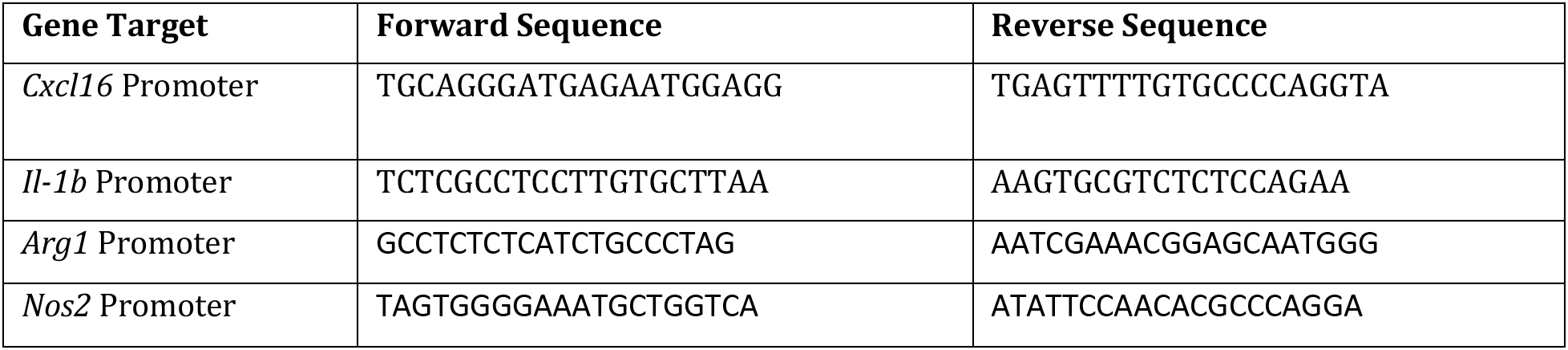
CUT & RUN qPCR Primer Sequences.

## Author Contributions

LM and AC conceived of the project and experimental design, analyzed data and wrote the manuscript. MT conducted and analyzed flow cytometry experiments. VB, JK and MR assisted with experiments and analysis. All authors contributed to the manuscript.

